# Trade-off between conservation of biological variation and batch effect removal in deep generative modeling for single-cell transcriptomics

**DOI:** 10.1101/2022.07.14.500036

**Authors:** Hui Li, Davis J. McCarthy, Heejung Shim, Susan Wei

**Affiliations:** School of Mathematics and Statistics, University of Melbourne, Melbourne, VIC 3010, Australia; Bioinformatics and Cellular Genomics, St Vincent’s Institute of Medical Research, Melbourne, VIC 3065, Australia; Melbourne Integrative Genomics, University of Melbourne, Melbourne, VIC 3010, Australia

**Keywords:** conservation of biological variation, batch effect, MINE, MMD, Pareto front, Pareto MTL, scRNA-seq, scVI

## Abstract

Single-cell RNA sequencing (scRNA-seq) technology has contributed significantly to diverse research areas in biology, from cancer to development. Since scRNA-seq data is high-dimensional, a common strategy is to learn low-dimensional latent representations better to understand overall structure in the data. In this work, we build upon scVI, a powerful deep generative model which can learn biologically meaningful latent representations, but which has limited explicit control of batch effects. Rather than prioritizing batch effect removal over conservation of biological variation, or vice versa, our goal is to provide a bird’s eye view of the trade-offs between these two conflicting objectives. Specifically, using the well established concept of Pareto front from economics and engineering, we seek to learn the entire trade-off curve between conservation of biological variation and removal of batch effects.

A multi-objective optimisation technique known as Pareto multi-task learning (Pareto MTL) is used to obtain the Pareto front between conservation of biological variation and batch effect removal. Our results indicate Pareto MTL can obtain a better Pareto front than the naive scalarization approach typically encountered in the literature. In addition, we propose to measure batch effect by applying a neural-network based estimator called Mutual Information Neural Estimation (MINE) and show benefits over the more standard Maximum Mean Discrepancy (MMD) measure. The Pareto front between conservation of biological variation and batch effect removal is a valuable tool for researchers in computational biology. Our results demonstrate the efficacy of applying Pareto MTL to estimate the Pareto front in conjunction with applying MINE to measure the batch effect.

## Background

Single-cell RNA sequencing (scRNA-seq) measures gene expression at single-cell resolution, allowing for the study of cell types, state, and trajectories to better understand heterogeneous cell states and dynamics in tissues, organs, and organism development. Since scRNA-seq data is high-dimensional and large-scale (e.g., gene expression of tens of thousands of genes for hundreds of thousands of cells or more), common analysis techniques often require discovering low-dimensional latent representations which capture underlying gene expression patterns in the high-dimensional data. Among widely used dimension reduction techniques (e.g., PCA [1], ZIFA [2], t-SNE [3], UMAP [4], PHATE [5]), methods based on deep neural network such as scVI [6] and SAUCIE [7] have emerged as powerful tools as they can be efficiently applied to the large-scale data.

While SAUCIE and scVI attempt to account for batch effects when learning the latent representations to ensure that the learned representations capture biological variations, these methods are not designed to learn the complex trade-off between conserving biological variation and removing batch effects. In this paper, building upon scVI, we aim to learn the trade-offs between these two conflicting objectives rather than prioritizing one over the other. Specifically, we borrow the concept of Pareto front from economics and engineering to construct the trade-off curve. We then use Pareto multi-task learning method [8] to estimate the Pareto front. Along the way, we introduce a new batch effect measure based on deep neural networks.

The rest of this section is as follows. We begin by reviewing the ZINB model for scRNA-seq data and the deep generative model proposed in scVI [6]. Then, we describe the concept of the Pareto front and summarise our contribution.

### ZINB model

The data resulting from an scRNA-seq experiment can be represented by an *n* × *G* matrix, ***x***, where each entry 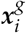 records the expression level measured for cell *i* and gene *g*. For each cell *i*, we observe a batch annotation *s_i_* ^1^. As the scRNA-seq data exhibits overdispersion and a large proportion of observed zeros, the zero-inflated negative binomial (ZINB) distribution [6] is commonly employed to model it. The ZINB model is a combination of negative binomial distribution for the expression count and logit distribution for the excessive zeros relative to the negative binomial distribution, perhaps due to failure in the assay reliably to capture information from genes with low expression. Specifically, conditional on batch ***s**_i_* and two latent variables ***z***_*i*_ and *l_i_*, we model 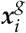 using the ZINB distribution. The latent variable ***z***_*i*_, a low-dimensional vector, potentially represents biological variation; its prior *p*(***z***) is 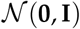. The latent variable *l_i_* represents log-library size; its prior *p*(*l*) is 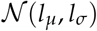, where *l_μ_*, *l_σ_* are set to be the empirical mean and variance of log-library size per batch.

Putting this altogether leads to the ZINB model:

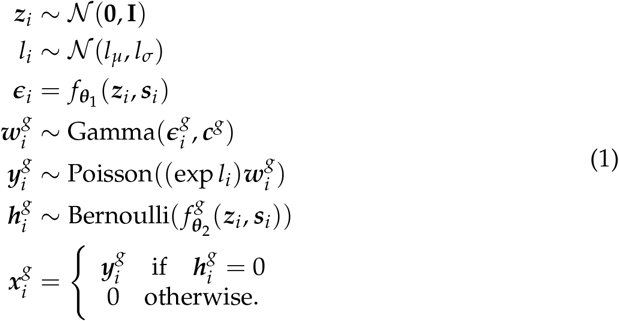

Above *f*_***θ***1_ and *f*_***θ***2_ are **decoder neural networks**. We shall denote by ***θ*** the concatenation of *θ*_1_ and *θ*_2_. The gene-specific inverse dispersion is denoted 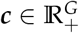 ^2^. As with 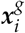, we use superscript notation to refer to a specific gene *g*. The notation 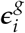 means the proportion of gene *g* expression in the whole expression of cell *i*, 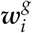 is the gene *g*’s expression proportion in cell *i* after gene inverse dispersion adjustment, 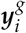g is the expression count of gene *g* in cell *i* from negative binomial distribution and 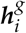 is the drop-out rate for gene *g* in cell *i* (thus 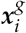 is zero-inflated negative binomial).

### Variational inference

The posterior distribution *p*(***z***, *l*|***x***)^3^ is unfortunately intractable. While we could employ MCMC to approximate the posterior, we shall instead consider the fast alternative of variational inference (VI). There are two main ingredients to VI: 1) an approximating family 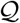, and 2) a criterion for determining the best member 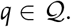. For the former, we turn to a mean-field variational family whereby each 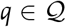 is stipulated to factor across the latent variables:

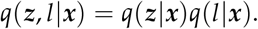

The distribution *q*(***z***|***x***) is further chosen to be multivariate Gaussian with diagonal covariance matrix and the distribution *q*(*l*|***x***) is chosen to be Gaussian with scalar mean and variance. The mean and variances of *q*(***z***|***x***) and *q*(*l*|***x***) will be learned using an **encoder neural network** applied to **x**. The collective weights of the encoder neural networks will be denoted by ***ϕ***.

For the second ingredient of VI, we adopt the conventional Kullback-Leibler (KL) divergence, i.e., we seek to minimize the KL divergence between *q*_***ϕ***_ (***z***, *l*|***x***) and the intractable true posterior *p*(***z***, *l* |***x***). It turns out that minimizing the KL divergence is equivalent to minimizing the negative evidence lower bound (ELBO), which we shall denote by *U_n_*(***ϕ***, ***θ***) (details in The loss function *U_n_* for the scVI generative model).

### Controlling batch effect

So far, there is nothing that prevents the learned variational distribution *q*_***ϕ***_ (***z***|***x***) from outputting latent representations ***z*** that are strongly correlated with the batch variable ***s***. In this work we set out to characterise the trade-off between learning ***z*** that is biologically meaningful and, simultaneously, disentangled from batch effects. For now, let *V_n_* (***ϕ***) be some measure of batch effect in the learned latent variable ***z***, where we take the convention that a lower value of *V_n_* is better, i.e., less batch effect. Note that, unlike the generative loss *U_n_*, the batch effect measure *V_n_* does not depend on the decoder parameter ***θ*** because the latent ***z*** only depends on the encoder parameter ***ϕ***.

Consider the bi-objective minimization problem,

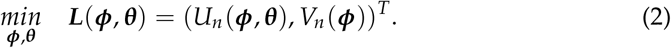

Note that this is a *vector* objective. Associated to this bi-objective problem is the so-called Pareto front:

#### Definition 1.

*Suppose we have a general optimisation problem with p objectives*:

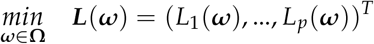

*where* **Ω** *is the parameter space and L_i_*: ***ω*** → ℝ, *i* = 1,…, *p. We say* ***ω*** ∈ **Ω** *is Pareto optimal if and only if it is non-dominated, i.e. there does not exist any*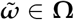 *such that* 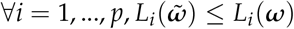 *with at least one strict inequality. The **Pareto front** is the set of all Pareto optimal points (Fig. 1A)*.

**Figure 1.**
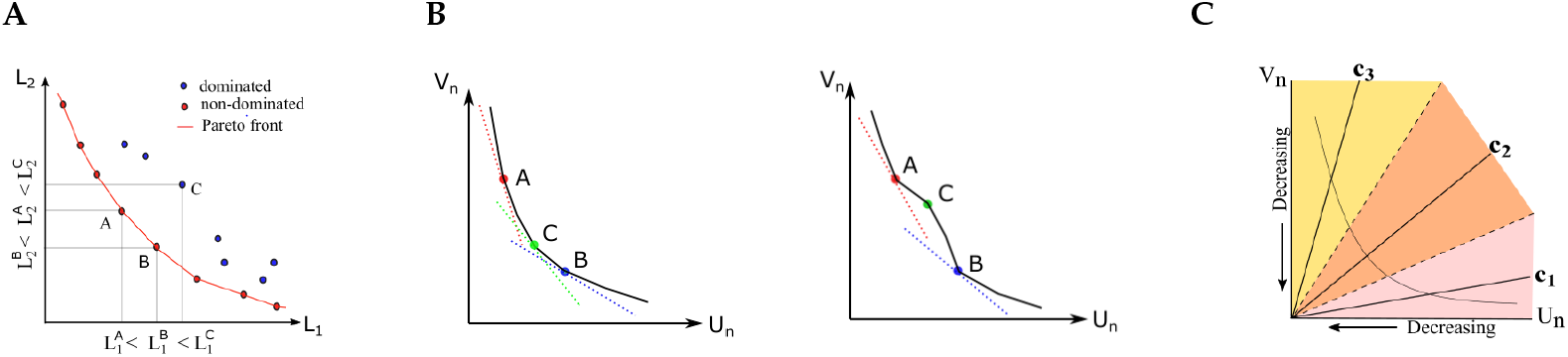
Panel A shows the Pareto front for an example bi-objective minimization problem. Point A and point B are non-dominated points on the Pareto front, while point C is dominated by both A and B. Panel B shows example Pareto candidates that can be discovered by the scalarization method for a convex (left) and a non-convex(right) Pareto front. In theory, scalarization cannot recover Pareto candidates in the non-convex part of a non-convex (right) Pareto front, such as Point C [9]. Panel C is a schematic of the Pareto MTL method [8], which first decomposes the bi-objective space into subregions according to a set of preference vectors *c_k_*, and then seeks a Pareto candidate within each subregion.

A naive strategy to obtain the Pareto front is via regularization,

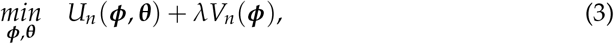

where λ ∈ ℝ can be viewed as a penalty factor. Under this strategy, Pareto candidates are generated by sweeping a list of penalty factors. Examples of this approach can be seen in [10] and [7]. In the former, to control batch effect, the generative loss of scVI is penalized by the Hilbert-Schmidt Independence Criterion (HSIC). In the latter, the generative loss of SAUCIE, a sparse autoencoder, is penalized by the Maximum Mean Discrepancy (MMD). We will later refer to these two methods as scVI+HSIC and SAUCIE+MMD respectively.

Now, the regularized objective in (3) is equivalent to the objective

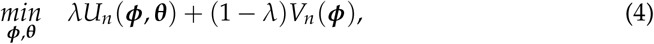

where *λ* ∈ [0,1]. This equivalent formulation can be recognized as the **scalarization scheme** in the multiobjective optimization field. In Fig. 1B, the scalarization approach for a given *λ* corresponds to one tangential point on the convex part of a Pareto front. If the true Pareto front is convex (left in Fig. 1B), the scalarization approach can in theory produce the full Pareto front (though it will remain challenging to find the proper set of *λ*’s). However, if the true Pareto front is non-convex as is often the case (right in Fig. 1B), the scalarization approach is not only inefficient but also unable to recover the Pareto optimal points on the non-convex parts, such as Point C [9].

### Contribution

In this research, we first improve upon the naive scalarization approach for estimating the Pareto front associated to (2) by applying the sophisticated Pareto multi-task learning (Pareto MTL) method proposed in [8]. We will see that with Pareto MTL we can find a set of “well-distributed” Pareto optimal points, in contrast to the scalarization approach which is typically only capabale of producing Pareto optimal points at the extremes of the Pareto front. Our second contribution is to propose a new batch effect measure based on the Mutual Information Neural Estimator (MINE) proposed in [11]. MINE leverages the expressiveness of deep neural networks to learn the mutual information (MI) between two variables, which in our case is the MI between the latent ***z*** and batch ***s***.

## Overview of methods

To allow readers to appreciate the results in the upcoming section, we briefly describe here the objectives *U_n_* and *V_n_* and how we performed Pareto front estimation.

### The loss function *U_n_* for the scVI generative model

The loss function *U_n_* associated to the scVI generative model [6] arises as follows. Minimizing the KL divergence^4^ between the variational distribution and the true posterior,

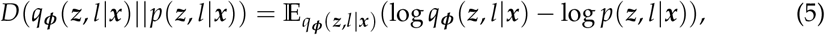

is equivalent to maximizing the so-called Evidence Lower Bound (ELBO),

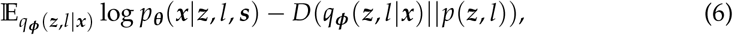

where *p*_***θ***_(***x***|***z***, *l*, ***s***) is the ZINB distribution defined in ZINB model. The prior *p*(***z***, *l*) in 6 is assumed to factor, i.e., *p*(***z***,*l*) = *p*(***z***)*p*(*l*). Then given a training set 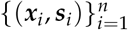, define

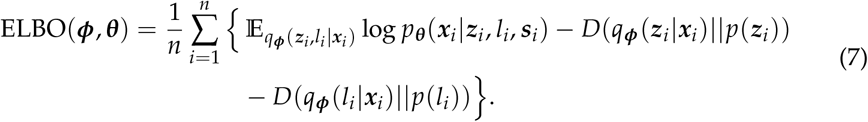

Throughout this paper we always employ *U_n_* (***ϕ***, ***θ***) = – ELBO(***ϕ***, ***θ***).

### Batch effect measure *V_n_* via MINE or MMD

In the Methods section, we will describe the details for each of the batch effect measures, MINE and MMD. It suffices for now to say that MINE can side-step the challenge of choosing a proper kernel bandwidth as is required in deploying MMD. MINE however incurs an additional computational cost because it has parameters of its own that need to be learned using adversarial training.

### Standardization of *U_n_* and *V_n_*

Standardization is an important precursor to the success of multi-objective optimization methods. Whether we use MINE or MMD for *V_n_*, we need to address the challenge posed by the highly imbalanced objectives *U_n_* and *V_n_*. To standardize *U_n_*, we minimize *U_n_*(***ϕ***, ***θ***) using minibatch stochastic gradient descent. Let *U_min_* and *U_max_* denote, respectively, the minimum and maximum value of generative loss observed across the mini-batches over all epochs. Similarly, to standardize *V_n_*, we first record *V_min_* and *V_max_*. The standardized objectives are then given by

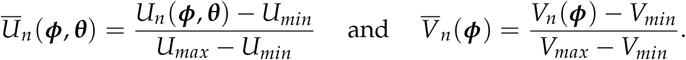

It must be noted that because their goal is not the explicit estimation of the Pareto front, neither scVI+HSIC nor SAUCIE+MMD perform standardization of *U_n_* and *V_n_* before performing the optimization in (3). But as our objective is to estimate the Pareto front, we must standardize each of *U_n_* and *V_n_*.

### Pareto front estimation

In all experiments in this work, we seek *K* = 10 Pareto front candidates. As we briefly outlined in the Background section, obtaining the Pareto front via the scalarization scheme is straightforward, entailing only a sweep of various *λ* ∈ [0,1] in Equation (4). Throughout, we employ *λ* ∈ {1/(*K* + 1),…, *K*/(*K* + 1)} for the scalarization scheme. We shall compare the naive scalarization approach to the more sophisticated Pareto MTL method (see Section Pareto MTL) for estimating the conservation of biological variation and batch effect removal Pareto front. We shall see that Pareto MTL produces Pareto candidates that are distributed in distinct regions of the trade-off curve rather than lumped in the extremes.

To obtain a complete Pareto front, we must add the two extreme points of the Pareto front corresponding to when only *U_n_* (***ϕ***, ***θ***) is minimized and when only *V_n_* (***ϕ***) is minimized. Note that when the *U_n_* and *V_n_* objectives are optimized on their own, no imbalance issue arises and hence no standardization is required. Finally, it is worth mentioning that in contrast to minimizing *U_n_* (***ϕ***, ***θ***), when *V_n_*(***ϕ***) is alone minimized, the output is simply some ***ϕ***_*T*_. To obtain the corresponding ***Θ***_*T*_, we then minimize *U_n_* (***ϕ***_*T*_, ***θ***) over *θ.*

## Results

For the bi-objective minimization problem in (2), we fix *U_n_* to be the loss function associated to the scVI generative model, while allowing for two possible batch effect measures *V_n_*. We will also consider two Pareto front estimation techniques. This makes for a total of four settings: 1) MINE or MMD for *V_n_* and 2) Pareto MTL or scalarization for Pareto front estimation. We begin by presenting the results of the best combination which is Pareto MTL with MINE.

### Pareto MTL with MINE

We first demonstrate that Pareto MTL with MINE can estimate a well-distributed Pareto front in the 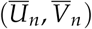 space. In Fig. 2A, in addition to the two extreme points, we plot the ten Pareto candidates produced by Pareto MTL for the Tabula Muris Marrow (TM-MARROW) dataset [12,13], a single cell transcriptome dataset from the model organism Mus musculus (see Single cell RNA-seq datasets). Ideally all 12 Pareto candidates should be non-dominated and appear on the Pareto front (see Definition 1). However, since stochastic optimization is not exact, we obtain dominated points. This explains why we display both the Pareto candidates along with the “culled” set of non-dominated points. We use point size to indicate the extent to which the generative loss is minimized. Points of smaller size stand for smaller generative loss minimization during training (i.e. preference vector closer to x-axis). The smallest and largest point size correspond to the extreme points. An analogous figure for the Macaque retina dataset (MACAQUE-RETINA) (see Single cell RNA-seq datasets) can be found in Fig S1 of Additional file 1.

**Figure 2.**
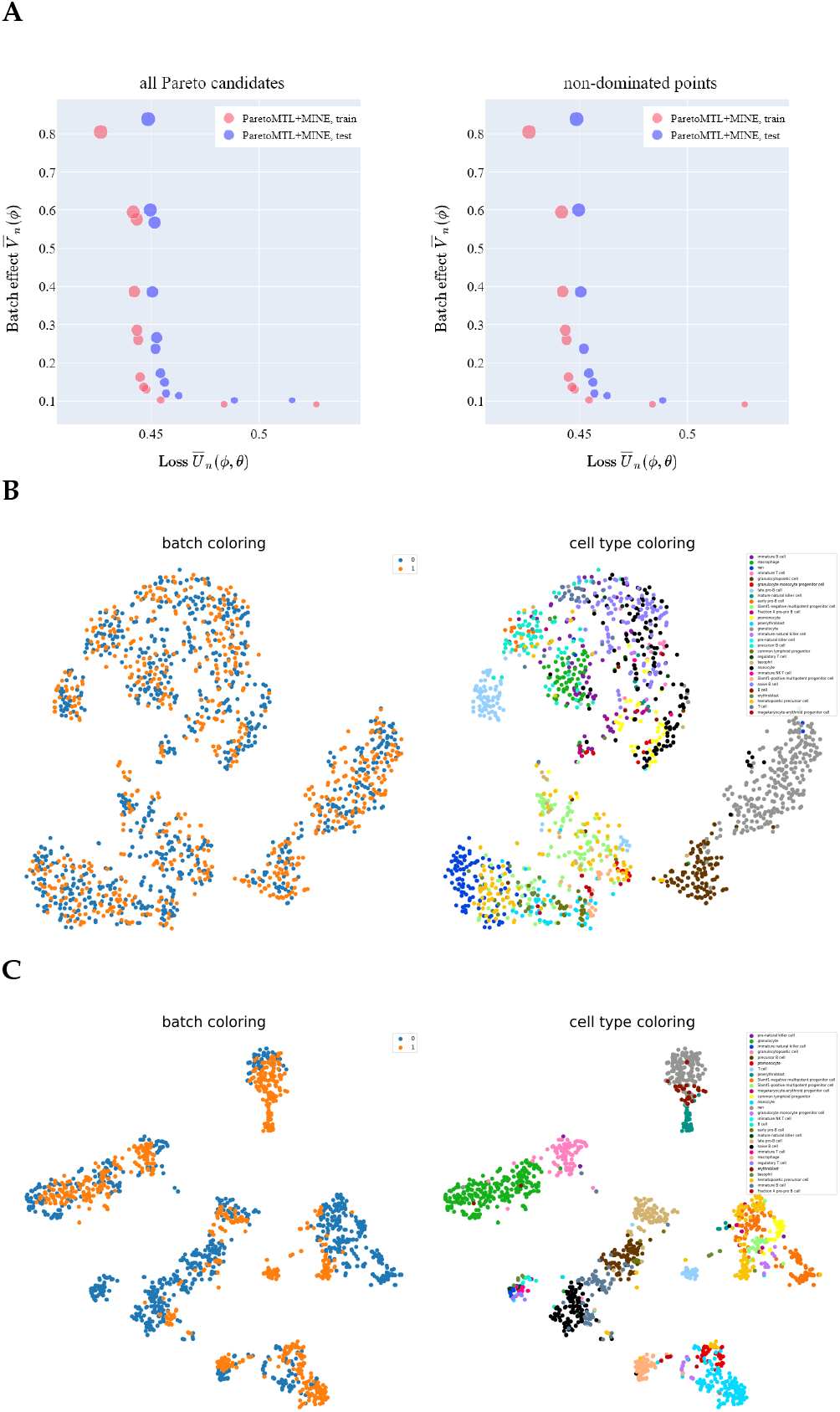
Pareto front in 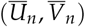 space estimated via Pareto MTL with MINE and associated t-SNE plots. Panel A shows all 12 Pareto candidates (left) and the culled candidates (right) in the 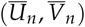 space on TM-MARROW dataset. Panel B and Panel C show the t-SNE plots of the latent ***z*** for the second and ninth subproblems on test set, respectively. Each point in the t-SNE plots indicates a cell and the points are colored by batches (left) and pre-annotated cell types (right). As expected, in Panel B we see less batch effect at the cost of poor clustering performance, while in Panel C, we see more batch effect but also better cell type clustering.

The trade-off points in the 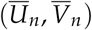 space have biological meaning. In Fig. 2B and Fig. 2C, we plot the latent ***z*** in a two dimensional plane via t-SNE method [3] for the second and ninth subproblems on the TM-MARROW test dataset, respectively. As expected for the second subproblem (Fig. 2B), more batch effect has been removed at the cost of less conservation of biological variation. In contrast, for the ninth subproblem (Fig. 2C), less batch effect has been removed with more conservation of biological variation. The result is consistent with the trade-offs in the left plot of Fig. 2A, where batch effect measure 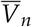 is lower (i.e. less batch effect) while generative loss 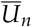 is higher (i.e. larger conservation of biological variation loss) for the second subproblem than for the ninth subproblem.

While t-SNE plots provide a useful visual aid to understand the various Pareto candidates, we can use surrogate metrics to obtain a quantitative understanding. Following [6], we use batch entropy (BE) to measure (roughly) how well latent ***z*** from different batches “mix”. We also use the following clustering metrics: Averaged Silhouette Width (ASW), Adjusted Rand Index (ARI), Normalized Mutual Information (NMI) and Unsupervised Clustering Accuracy (UCA). For all the above surrogate measures, higher is better. In particular, higher BE means better “mixing” of latent ***z*** from different batches. Higher ASW means clusters of latent ***z*** are further apart. ARI, NMI and UCA measure how consistent cell clustering of latent ***z*** by K-Means is with cell clustering based on the pre-annotated cell types and higher values correspond with better clustering. As expected, the second subproblem (Fig. 2B) has larger BE and smaller clustering surrogate metrics with

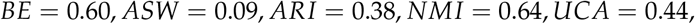

compared with the ninth subproblem (Fig. 2C) where

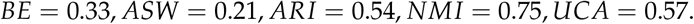

### Pareto MTL versus scalarization

Here we present results comparing Pareto MTL (see Pareto MTL) and the naive scalarization approach for estimating the 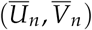 Pareto front when *V_n_* is given by either the MINE or MMD measure. The results on the TM-MARROW dataset are shown in Fig. 3; results on the MACAQUE-RETINA dataset can be found in Additional file 1 Fig S2. Points of smaller size stand for smaller generative loss minimization during training (i.e. smaller *λ* for scalarization). Note that the two extreme points are the same for Pareto MTL and scalarization.

**Figure 3.**
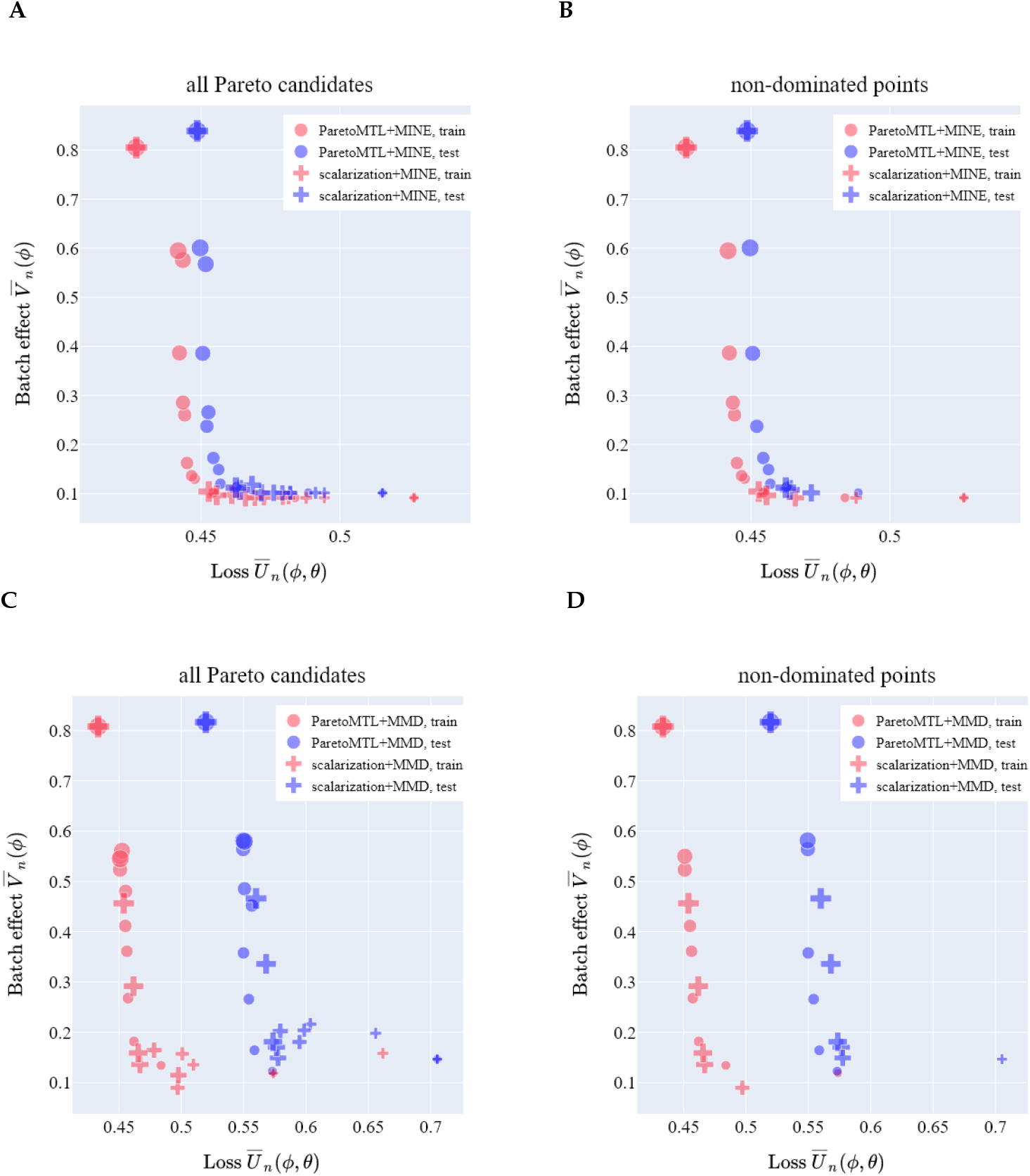
Pareto MTL vs scalarization. In Panels A and C, we show the 12 Pareto candidates produced when MINE and MMD, respectively, are used to measure the effect in the TM-MARROW dataset. In the corresponding Panels B and D, we show the non-dominated points only. When batch effect is measured using MINE, Pareto MTL produces a more diverse set of trade-offs, while scalarization tends to produce trade-offs at the extreme regions. When MMD is the batch effect indication, Pareto MTL seems to perform similarly to scalarization.

When MINE is the batch effect measure, we observe that as we progress from subproblem 1 to 10, the generative loss 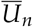 decreases while the batch effect measure 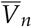 increases(Fig. 3A, Additional file 1 Fig S2). We describe in this case the subproblem ordering is well respected. Furthermore, we see that Pareto MTL is better than the scalarization scheme for estimating a well-distributed Pareto front (Fig. 3B, Additional file 1 Fig S2).

When MMD is the batch effect measure, we first notice that the subproblem ordering is not well-respected by neither Pareto MTL nor the scalarization scheme (Fig. 3C). We suspect this issue arises from the difficulty in selecting a proper MMD bandwidth across the subproblems. Second, Pareto MTL performs either similarly (Fig. 3D) or better (Additional file 1 Fig S2) than scalarization for estimating an well-distributed Pareto front.

The visual comparison in Fig. 3 and Fig S2 is based on *one* random splitting of the dataset into training and testing set. In order to compare Pareto MTL and the scalarization scheme more systematically, we focus on three metrics (for which higher is better) that can be measured on Pareto fronts:

- *percentage* of non-dominated points in all Pareto candidates
- *hypervolume* [14] evaluates the coverage area of the estimated Pareto front (see Hypervolume)
- *number of distinct choices* (NDC) [15] measures the number of meaningful Pareto solutions that are sufficiently distinct to one another (see NDC)

These three metrics evaluate three respective properties of the Pareto front:

- *cardinality*, which quantifies the number of non-dominated points,
- *convergence*, which evaluates how close a set of estimated non-dominated points is from the true Pareto front in the objective space, and
- *distribution*, which measures how well distributed the points are on the estimated Pareto front [16].

The Pareto MTL and scalarization scheme (either with MINE or MMD as batch effect indication) are each run for 10 training-testing splits. The percentage, hypervolume and NDC of the Pareto fronts in the 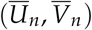 space on the two datasets (see Single cell RNA-seq datasets), are shown in Table 1.

**Table 1.**
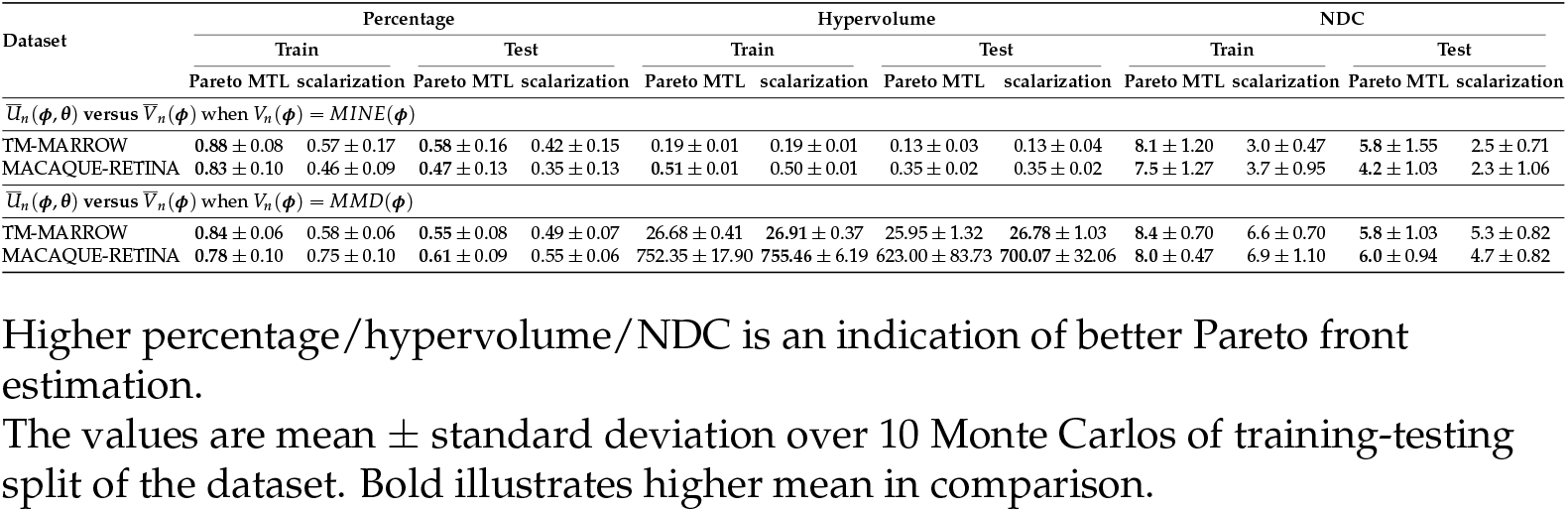
Pareto MTL versus scalarization.

We observe higher percentages of non-dominated points for Pareto MTL than scalarization. In terms of hypervolume, Pareto MTL and scalarization perform similarly for both MINE and MMD. Note that hypervolume for Pareto MTL with MMD is not significantly higher than scalarization with MMD when considering the standard deviation. One reason could be that the relative values of the hypervolume metric of two non-dominated point sets (i.e. which set has a larger hypervolume and which set has a smaller hypervolume) depend on the chosen reference point [16]. As for NDC, Pareto MTL with MINE is significantly better than scalarization with MINE; it is consistent with the observation that Pareto MTL with MINE can estimate a more diverse Pareto front than scalarization with MINE (Fig 3B). Pareto MTL with MMD also produces Pareto fronts with higher NDC than scalarization with MMD, but the superiority is not as dramatic as that between Pareto MTL with MINE and scalarization with MINE.

### Pareto MTL with MINE versus Pareto MTL with MMD

Though both MINE and MMD measure batch effect, Pareto MTL with MINE and Pareto MTL with MMD cannot be directly compared in the 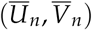 space. Specifically, MINE measures the KL divergence between distributions (see see MINE for measuring batch effect *V_n_*) while MMD is the maximum mean embedding distance between distributions (see MMD for measuring batch effect *V_n_*). We can however make meaningful comparisons by introducing surrogate measures for *V_n_*.

We use two surrogate measures for *V_n_*: 1) the aforementioned batch entropy (BE) and 2) nearest neighbor (NN). NN here refers to a method proposed in [17] to estimate the mutual information (MI) between the continuous latent ***z*** and discrete batch ***s*** instead of the traditional k-nearest neighbor algorithm for classification. Larger values of BE correspond to smaller batch effect, while larger values of NN correspond to higher batch effect. In Fig S4 and Fig S5, see Additional file 1, we show simulation results that indicate NN estimates well the mutual information between a continuous variable and a categorical variable. We should note that NN cannot be used during neural network training because it is not amenable to back-propagation.

We could also use the aforementioned clustering metrics (ASW, ARI, NMI and UCA) as the evaluation surrogates for *U_n_* since they are more interpretable than the ELBO. Therefore, in total we can produce ten types of trade-off curves for evaluation purposes, i.e. five measures of 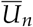 (*U_n_* itself and four clustering surrogates) cross two surrogate measures of 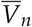 (BE and NN). For brevity, we will only present a subset of five to compare Pareto MTL with MINE and Pareto MTL with MMD: *U_n_* versus *NN*, and negative ASW/NMI/ARI/UCA versus negative BE. The trade-offs of *U_n_* versus *NN*, negative ASW/NMI versus negative BE for TM-MARROW dataset are shown in Fig. 4 and the trade-offs of negative ARI/UCA versus negative BE for TM-MARROW dataset in Additional file 1 Fig S6.

**Figure 4.**
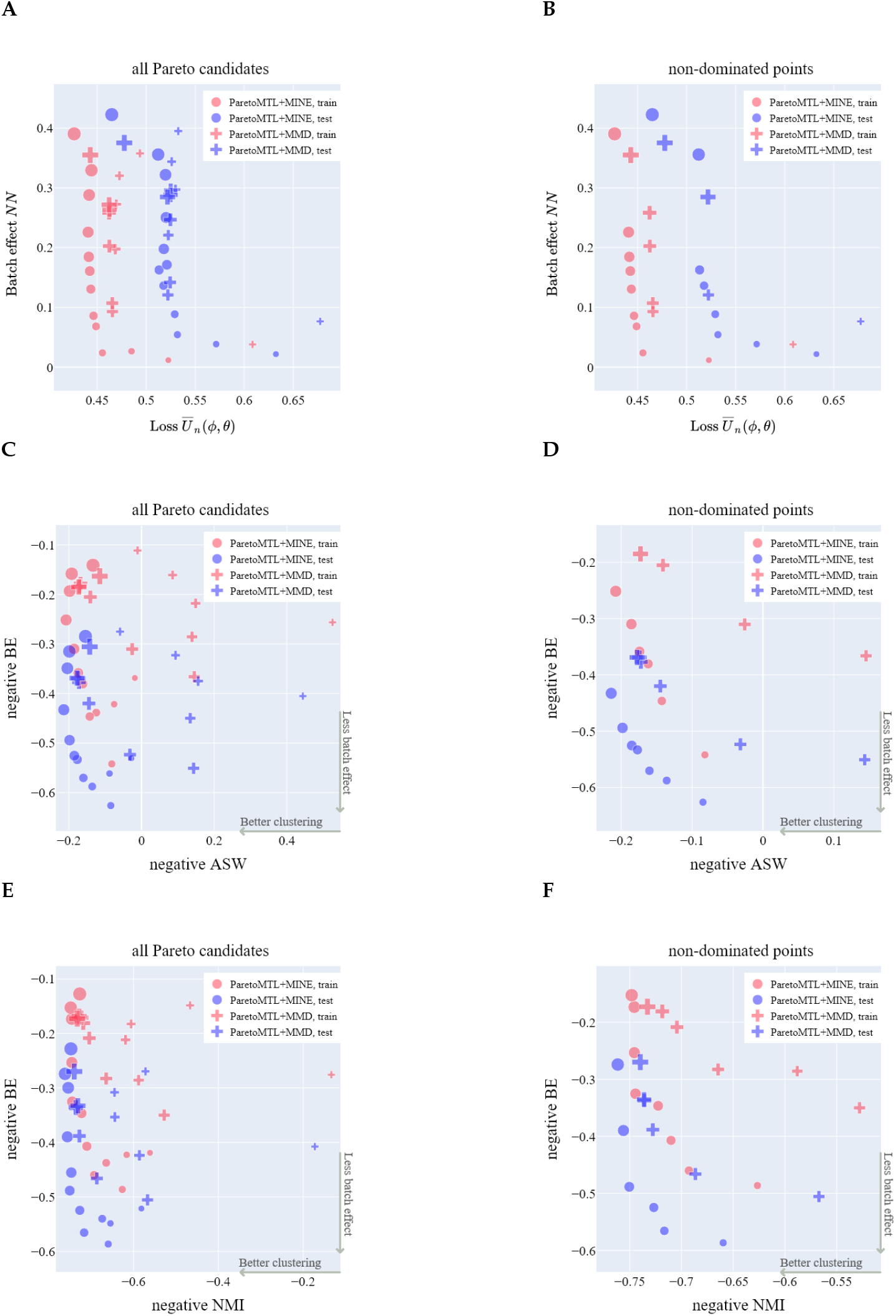
The trade-off curves of surrogate metrics. Compare Pareto MTL with MINE and Pareto MTL with MMD to estimate three types of trade-offs of surrogate metrics for all Pareto candidates and non-dominated points on the TM-MARROW dataset: (A,B) trade-offs between generative loss *U_n_* and mutual information estimator *NN*; (C,D) the trade-offs between negative ASW and negative BE; (E,F) the trade-offs between negative NMI and negative BE

The trade-offs in the 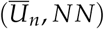, *NN*) space for Pareto MTL with MINE (Fig. 4A) respects the subproblem ordering as in the 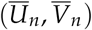 space in Fig. 3A. In contrast, Pareto MTL with MMD produces trade-offs in the (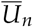, *NN*) space in disarray (Fig. 4A). Besides, we observe that Pareto MTL with MINE estimates a better set of non-dominated (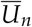, *NN*) trade-off points which dominates that estimated by Pareto MTL with MMD (Fig. 4B).

Similarly, in the negative ASW/NMI and negative BE space, Pareto MTL with MINE produces trade-offs that respect subproblem ordering better than Pareto MTL with MMD does (Figs. 4C, 4E). With the exception of some points at the extremes, Pareto MTL with MINE has a clearer trend than Pareto MTL with MMD, in the sense that as negative clustering metrics decrease, negative BE increases. Meanwhile, Pareto MTL with MINE produces a set of non-dominated trade-off points which dominates those from Pareto MTL with MMD (Figs. 4D and 4F). Similar conclusion can be drawn for trade-offs between negative ARI/UCA and negative BE (Additional file 1 Fig S6).

Analogously we could obtain trade-offs in the surrogate space for scalarization with MINE and scalarization with MMD (Additional file 1 Fig S7). The results provide extra support that MINE is better than MMD for measuring batch effect since scalarization with MINE obtains better trade-offs in terms of subproblem ordering of the Pareto candidates and convergence of the non-dominated points. However, as already seen in Fig. 3B, the problem for scalarization with MINE is that it can only recover a small part of the Pareto front (i.e. diversity problem).

Finally, we can quantitatively compare Pareto MTL with MINE and Pareto MTL with MMD by again evaluating percentage, hypervolume and NDC, but this time in the surrogate bi-objective space. The results for 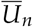 versus *NN* and negative ASW/NMI versus negative BE on the two datasets are shown in Table 2; the results for negative ARI/UCA versus negative BE are shown in Additional file 1 Table S1. Compared with Pareto MTL with MMD, Pareto MTL with MINE produces better surrogate Pareto fronts with 1) higher or similar percentage of non-dominated points (i.e. higher or similar cardinality), 2) larger hypervolume (i.e. better convergence) and 3) higher or similar NDC (i.e. better or similar diversity).

**Table 2.** Pareto MTL with MINE versus Pareto MTL with MMD.

| Dataset | Percentage |  |  |  | Hypervolume |  |  |  | NDC |  |  |  |
| --- | --- | --- | --- | --- | --- | --- | --- | --- | --- | --- | --- | --- |
|  | Train |  | Test |  | Train |  | Test |  | Train |  | Test |  |
|  | MINE | MMD | MINE | MMD | MINE | MMD | MINE | MMD | MINE | MMD | MINE | MMD |
| $\bar{U}_n(\phi, \theta)$ versus $NN$ | | | | | | | | | | | | |
| TM-MARROW | 0.85 ± 0.07 | 0.48 ± 0.07 | 0.58 ± 0.18 | 0.35 ± 0.12 | 4.35 ± 0.04 | 3.91 ± 0.16 | 4.29 ± 0.06 | 3.85 ± 0.21 | 7.7 ± 0.82 | 5.4 ± 0.52 | 5.6 ± 1.35 | 4.1 ± 1.20 |
| MACAQUE-RETINA | 0.72 ± 0.10 | 0.27 ± 0.08 | 0.47 ± 0.13 | 0.14 ± 0.04 | **293.82 ± 1.77** | 273.48 ± 4.23 | **295.15 ± 4.17** | 257.92 ± 8.90 | 4.1 ± 0.74 | 2.0 ± 0.00 | 3.8 ± 1.55 | 1.2 ± 0.42 |
| -ASW versus -BE |  |  |  |  |  |  |  |  |  |  |  |  |
| TM-MARROW | 0.45 ± 0.06 | 0.43 ± 0.09 | 0.48 ± 0.10 | 0.41 ± 0.05 | 0.29 ± 0.02 | 0.16 ± 0.02 | 0.38 ± 0.01 | 0.30 ± 0.02 | 4.6 ± 0.84 | 4.2 ± 0.79 | 4.9 ± 1.37 | 3.7 ± 0.48 |
| MACAQUE-RETINA | 0.61 ± 0.07 | 0.33 ± 0.10 | 0.64 ± 0.10 | 0.34 ± 0.11 | 0.32 ± 0.01 | 0.15 ± 0.02 | 0.30 ± 0.01 | 0.19 ± 0.03 | 4.4 ± 0.84 | 3.1 ± 0.57 | 5.5 ± 0.85 | 2.6 ± 0.70 |
| -NMI versus -BE |  |  |  |  |  |  |  |  |  |  |  |  |
| TM-MARROW | 0.56 ± 0.08 | 0.49 ± 0.08 | 0.58 ± 0.12 | 0.40 ± 0.09 | 0.28 ± 0.01 | 0.16 ± 0.02 | 0.37 ± 0.01 | 0.31 ± 0.02 | 6.0 ± 0.67 | 4.5 ± 0.85 | 5.0 ± 1.33 | 4.3 ± 0.95 |
| MACAQUE-RETINA | 0.38 ± 0.11 | 0.23 ± 0.11 | 0.49 ± 0.18 | 0.33 ± 0.12 | 0.26 ± 0.02 | 0.11 ± 0.02 | 0.25 ± 0.01 | 0.14 ± 0.02 | 3.5 ± 0.85 | 2.5 ± 0.85 | 4.5 ± 1.27 | 3.5 ± 1.08 |

The Pareto fronts—generative loss $U_n$ versus mutual information estimator $NN$ and negative ASW/NMI versus negative BE from Pareto MTL with MINE and Pareto MTL with MMD are compared.
Better Pareto front has higher percentage/hypervolume/NDC.
The values are mean ± standard deviation over 10 Monte Carlos of training-testing split of the dataset. Bold illustrates higher mean in comparison.

## Discussion

We briefly discuss limitations and future work in this section. Although we have shown MINE to be superior to MMD in the current context of the work, MINE is computationally more demanding than MMD. Furthermore, using MINE in conjunction with scVI requires adversarial training which can be unstable in the hands of an unpracticed user. Finally, it is possible that designing the proper neural network architecture in MINE is a difficult task, though we did not encounter this in our data analysis. In fact, based on the analysis we have performed so far, the trade-off curves seem quite robust to the MINE architecture in contrast to the high sensitivity around MMD kernel and bandwidth choice.

In future work, we aim to incorporate other deep generative models into our framework. Though we have focused in this work on the scVI generative model for scRNA-seq data, our research is broadly applicable to other generative models for scRNA-seq data. For instance Pareto MTL with MINE can be wrapped around such deep generative models as SAUCIE and DESC [18]. Future work may also consider generalizing to other nuisance factors such as quality control metrics which assess the errors and corrections of aligning transcripts to some reference genome during scRNA sequencing.

## Conclusion

We propose using Pareto MTL for estimation of Pareto front in conjunction with MINE for measurement of batch effect to produce the trade-off curve between conservation of biological variation and removal of batch effect. We first demonstrated that Pareto MTL improves upon the naive scalarization approach in finding the Pareto front. In particularly, Pareto MTL produces well-distributed trade-off points in contrast to the scalarization approach which produces points in the extremes. We next demonstrated that the new batch effect measure based on MINE is preferable to the more standard MMD measure in the sense that the former produces trade-off points that respect subproblem ordering and are interpretable in surrogate metric spaces.

## Methods

### MINE for measuring batch effect *V_n_*

Mutual information *I*(***z***, ***s***) measures the dependence between continuous latent variable ***z*** and categorical ***s*** as follows

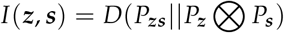

where *P*_***zs***_ is the joint distribution and *P****z*** ⊗ *P*_***s***_ is the product of the two marginal distributions. It was shown in [11] that we can estimate *I*(***z***, ***s***) by maximizing a lower bound *I*_ψ_(***z***, ***s***) defined as

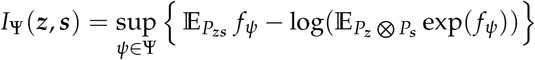

where *f*_ψ_ : ℝ^*d_z_*^ × {0,1}^*B*^ → ℝ is a deep neural network with parameters *ψ* ∈ Ψ.

Let ***z***_*i*_ be a realization from the posterior distribution *q*_***φ***_(***z***|***x**_i_*). We should point out that though (***z***_*i*_, ***s***_*i*_) are independent across *i*, they are not identically distributed since ***x***_*i*_ varies. We will be considering the so-called aggregated variational posterior for latent ***z***, see discussion in [10], which in our case is a *n*-component Gaussian mixture with equal weights. We shall treat 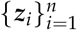 as a sample from this Gaussian mixture and ***s***_*i*_ is the known coupled batch for each ***z***_*i*_. To obtain a sample from the marginal *P*_***s***_ we shuffle ***s***_*i*_ to break the coupling with ***z***_*i*_; we denote this sample 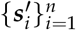. This leads us to define the following measure for batch effect:

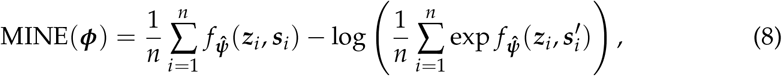

where 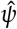 is the result of using minibatch stochastic gradient ascent, see Alg. 1.

The simulation studies (Additional file 1 Fig S4, Fig S5) demonstrate that MINE approximates well the true MI between a categorical variable and a continuous variable.

#### Algorithm 1 Train MINE

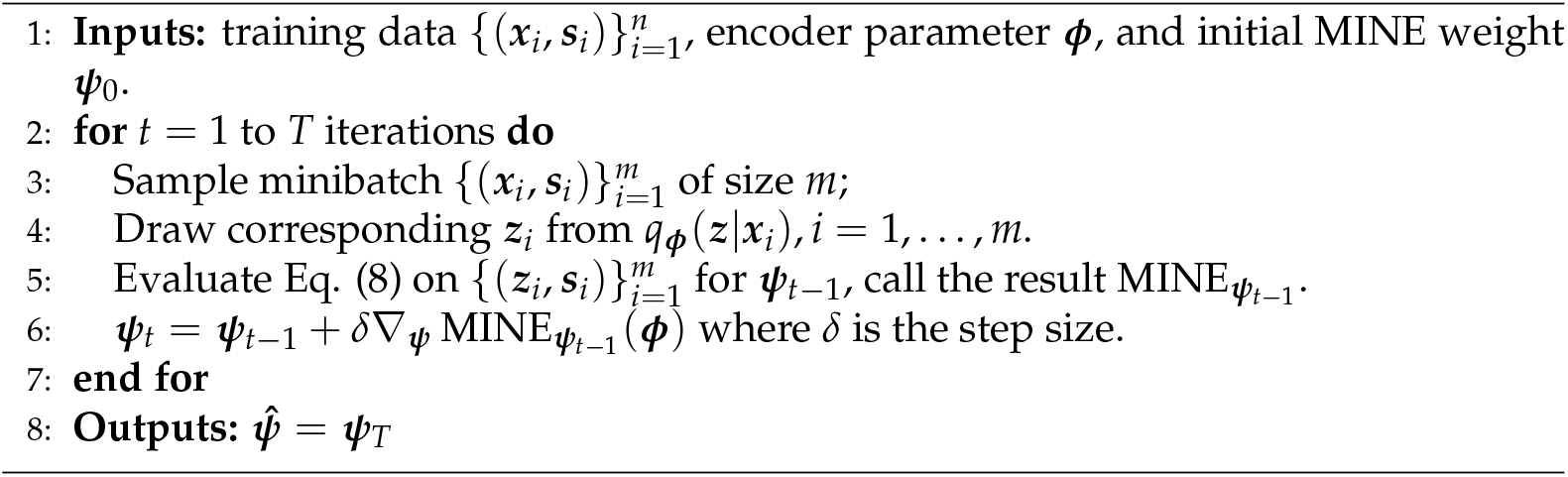

### MMD for measuring batch effect *V_n_*

The maximum mean discrepancy (MMD) [19] measures the discrepancy between two densities *p* and *q* using the unit ball in a reproducing kernel Hilbert space 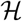 with associated kernel *k*(·, ·). The squared MMD is given by

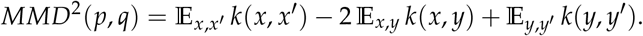

where *x* and *x*′ are independent random variables with distribution *p* and *y* and *y*′ are independent random variables with distribution *q*.

For simplicity let us assume we have two batches. To measure the batch effect, we will apply MMD to measure the disparity between the distributions ***z***|***s*** = 0 and 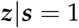. Let us denote 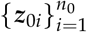 for the samples from batch 0 and 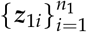 for the samples from batch 1. We estimate the MMD between 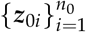 and 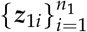 using the biased estimator

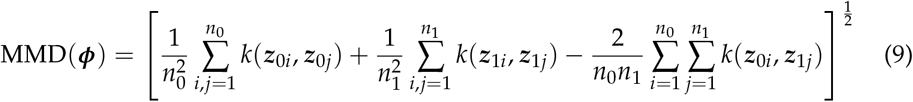

For development of this estimator see Section 2.2 in [19]. As in the previous section, the latent variable ***z***_*i*_ is sampled from *q*_***ϕ***_(***z***|***x***_*i*_) though in the right hand side of Eq. (9) we have suppressed the dependence on ***ϕ***.

Following [10], we shall take the kernel *k* in our experiments to be a mixture of 5 Radial Gaussian kernels:

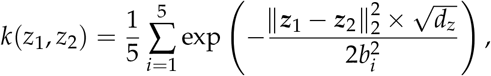

where ***z***_1_ and ***z***_2_ are any pair of latent ***z***, (*b*_1_,…, *b*_5_) is the chosen bandwidths and *d*_***z***_ is the dimension of ***z***. To avoid different scales for each dimension, the latent ***z*** is standardized such that each dimension has zero mean and unit standard deviation.

### Pareto MTL

Here we describe the application of the Pareto MTL method proposed in [8] to our work. Throughout our experiments, we will use the set of preference vectors

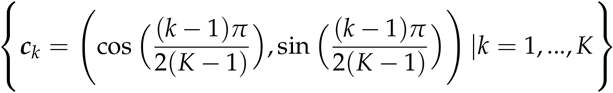

which is sensible since our particular choices of *U_n_* and *V_n_* will always be non-negative. Associated to the *k^th^* preference vector ***c***_*k*_ is the constrained optimisation subproblem

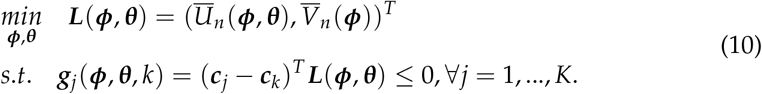

To begin, we seek initialization of ***ψ*** and ***θ*** such that the bi-objective ***L***(***ψ**, **θ***) starts its minimization from a place near the preference vector ***c***_*k*_. The initialization proceeds as follows.

1. Obtain the Lagrange multipliers *β_j_* by solving:

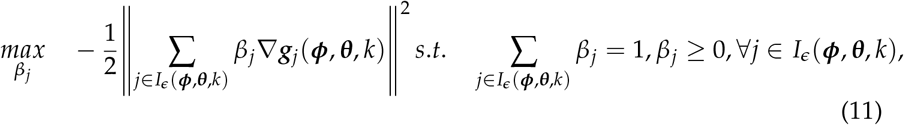

where *I_ϵ_*(***ψ**, **θ**, k*) = {*j*|*g_j_*(***ψ***, ***θ***, *k*) ≥ – *ϵ, j* = 1,…, *K*} and *ϵ* is a pre-defined small positive value.
2. Update (***ψ, θ***) by descending the gradient vector ***d*** with step size ***δ***:

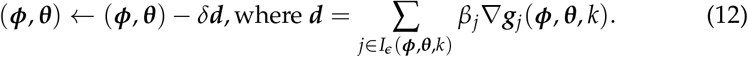

After initialization, the subproblem in Eq. (10) is solved as another dual problem:

1. Obtain the Lagrange multipliers *λ*_1_, *λ*_2_ and *β_j_* by solving:

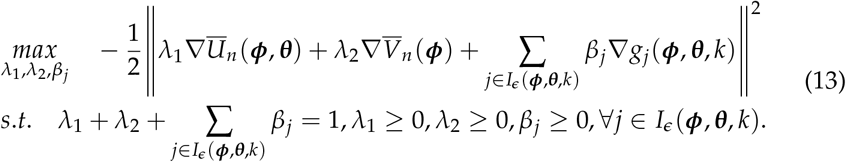
2. Update (***ψ, θ***) by descending the gradient vector ***d*** with step size *δ*:

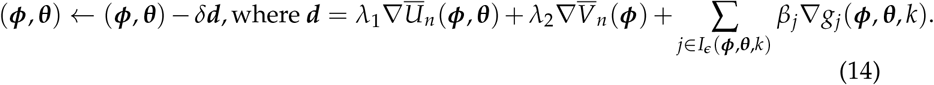

### Pareto MTL with MINE

When 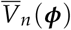 is based on MMD(***ψ***), the implementation of Pareto MTL is straightforward. However, when 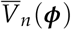 is based on MINE(***ψ***), we employ adversarial training since the MINE network has its own parameters ***ψ*** that need to be learned. Specifically, MINE(***ψ***) is **maximized** over the MINE neural network parameters ***ψ*** before every **minimization** step over the variational autoencoder parameters ***ψ*** and ***θ***. Alg. 3 gives an overview of Pareto MTL with MINE via adversarial training for the *k*-th subproblem. An estimate of the Pareto front can be obtained by running Alg. 3 for *k* = 1,…, *K*.

#### Algorithm 2 Initialization for Pareto MTL

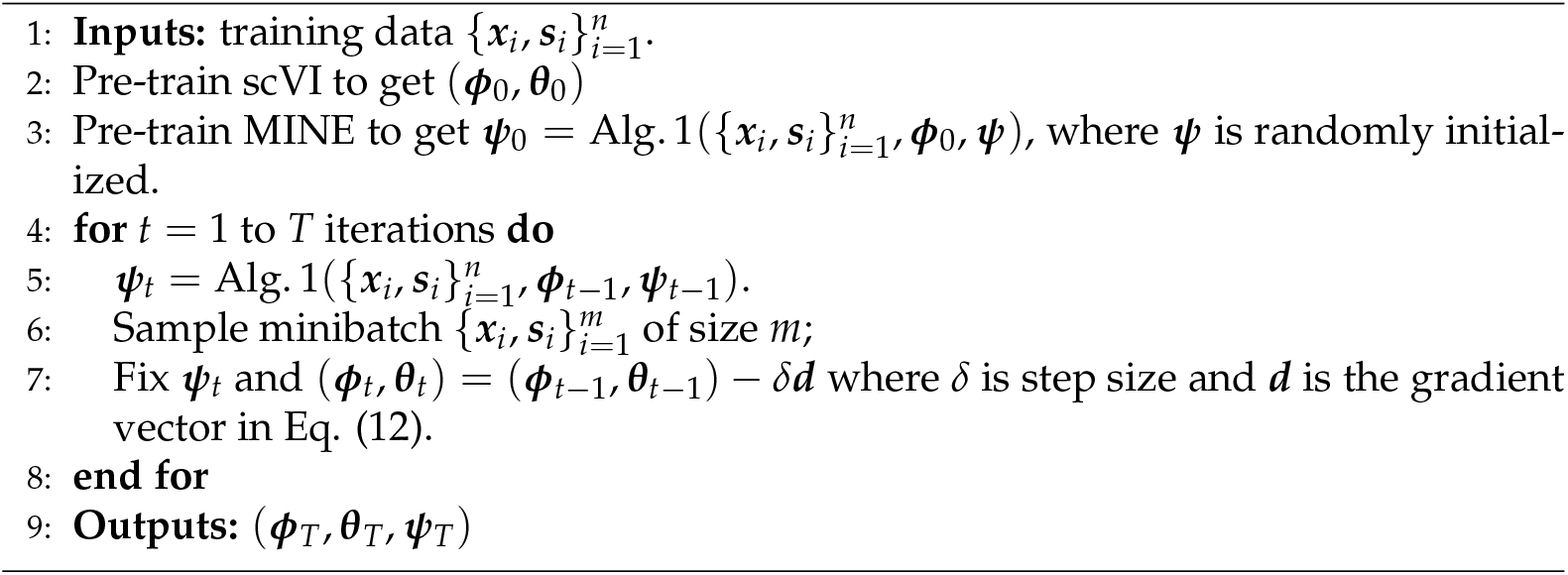

#### Algorithm 3 Pareto MTL with MINE

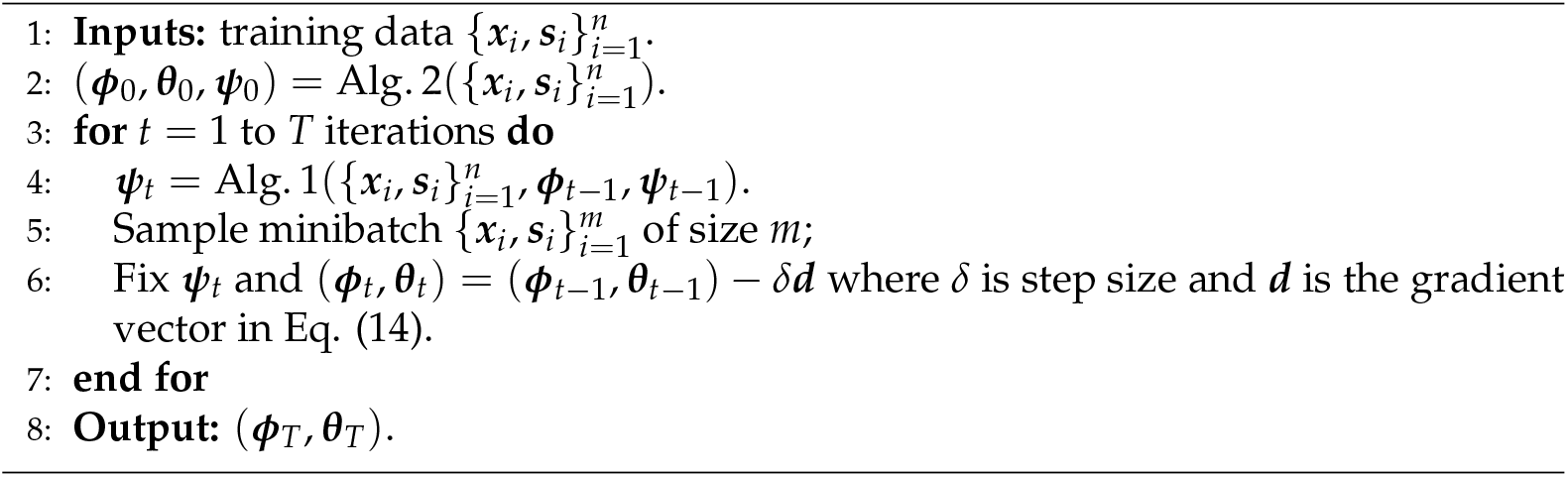

### Plotting the 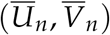 Pareto fronts

For each of the subproblems in Pareto MTL or scalarization, a resulting encoder-decoder pair is produced (***ϕ***_*T*_, ***θ***_*T*_). To understand such figures as Figs. 2A and 3, we describe how we evaluated *U_n_* and *V_n_* given a pair (***ϕ***_*T*_, ***θ***_*T*_). It is straightforward to evaluate *U_n_* given (***ϕ***_*T*_, ***θ***_*T*_).

When MMD is used to measure batch effect, it is also easy to evaluate *V_n_* given ***ϕ***_*T*_. However, when MINE is used to measure batch effect, the evaluation of *V_n_* is more subtle since MINE has its own training parameters unlike MMD. We first train a de-novo MINE neural network to its optimal (***ψ*** = ***ψ***\*, see MINE for measuring batch effect *V_n_*) with the encoder weight ***ϕ*** fixed at ***ϕ***_*T*_. Then with ***ψ*** = ***ψ***\*, Figs. 2A and 3 proceed to plot *V_n_* = MINE(***ϕ***_*T*_).

### Training details

To estimate the expectation in the ELBO, a single ***z*** is sampled from the variational distribution *q*_***ϕ***_(***z***_*i*_|***x***_*i*_) and likewise for *l* from *q*_***ϕ***_(*l_i_*|***x***_*i*_). When possible, we use analytic expressions for the KL divergences between two Gaussian distributions.

We use the same encoder and decoder architectures as the scVI in [6]. For MINE neural network architecture, we use 10 fully connected layers with 128 hidden nodes and ELU activation for each layer. The weights ***ψ*** of the MINE neural network is initialized by a normal distribution.

The hyperparameters for Pareot MTL on the TM-MARROW and MACAQUE-RETINA datasets are shown in Table 3. We employ the same hyperparameters for scalarization as its Pareto MTL counterpart. We use Adam optimiser (a first-order stochastic optimizer) with the parameter *ϵ* = 0.01 which improves the numerical stability of the optimizer and other parameters at their default values. The batch size for TM-MARROW and MACAQUE-RETINA are 128 and 256 respectively.

**Table 3.**
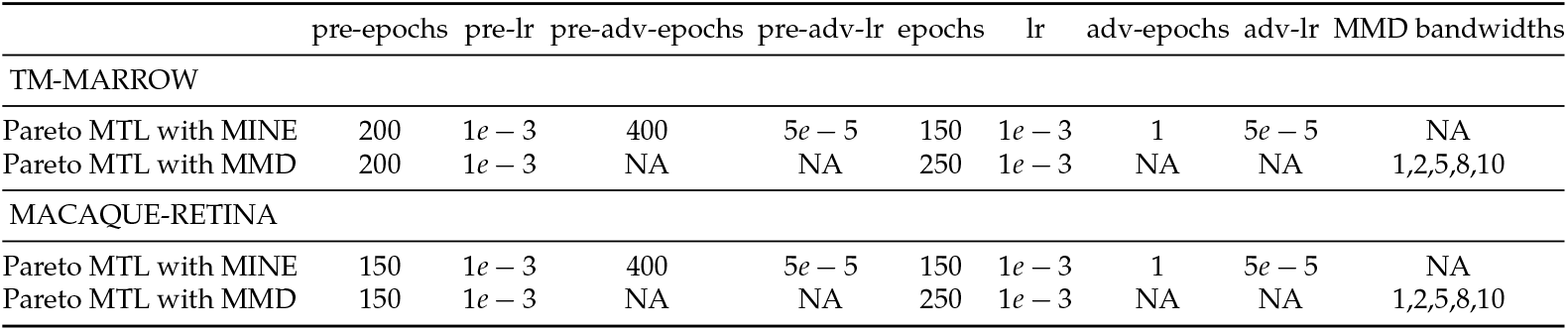
Hyperparameters for Pareto MTL. The abbreviation “pre” means pre-training, “adv” means MINE adversarial training, “lr” means learning rate.

|  | pre-epochs | pre-lr | pre-adv-epochs | pre-adv-lr | epochs | lr | adv-epochs | adv-lr | MMD bandwidths |
| --- | --- | --- | --- | --- | --- | --- | --- | --- | --- |
| TM-MARROW |  |  |  |  |  |  |  |  |  |
| Pareto MTL with MINE | 200 | $1e-3$ | 400 | $5e-5$ | 150 | $1e-3$ | 1 | $5e-5$ | NA |
| Pareto MTL with MMD | 200 | $1e-3$ | NA | NA | 250 | $1e-3$ | NA | NA | 1,2,5,8,10 |
| MACAQUE-RETINA |  |  |  |  |  |  |  |  |  |
| Pareto MTL with MINE | 150 | $1e-3$ | 400 | $5e-5$ | 150 | $1e-3$ | 1 | $5e-5$ | NA |
| Pareto MTL with MMD | 150 | $1e-3$ | NA | NA | 250 | $1e-3$ | NA | NA | 1,2,5,8,10 |

### Evaluating the estimated Pareto front

#### Hypervolume

In Additional file 1 Fig S3, we illustrate the hypervolume of a Pareto front. First, a reference point is chosen which has larger value in at least one dimension with no smaller values in all other dimensions than all the estimated points. Then each estimated point forms a rectangle with the reference point. The area of the union of all these rectangles is the hypervolume. Note that the reference point is shared among the methods to allow for proper comparison of Pareto front approximations.

#### NDC

In Additional file 1 Fig S3, we demonstrate how to obtain NDC of a Pareto front. For simplicity, in a set of Pareto candidates, suppose the range between maximum and minimum values of the loss *L*_1_ and the similar range for the loss *L*_2_ are both divisible by a pre-specified value *μ*. Then the ranges for the two losses are divided into a grid of squares with width *μ*. Each square is an indifference region. If there is at least one non-dominated point in the indifference region, the NDC for that region is one, otherwise 0. The final NDC is the sum of NDC for each indifference region. Note that the grids of squares are shared among the methods to allow for proper comparison of Pareto front approximations.

### Single cell RNA-seq datasets

We examined two datasets to evaluate the efficacy of Pareto MTL with MINE for Pareto front estimation. The TM-MARROW dataset is an integration of two bone marrow datasets (MarrowTM-10x, MarrowTM-ss2) from the Tabula Muris project [12,13]. The read counts in MarrowTM-ss2 are first standardized by mouse gene length and only cells with total count larger than 100 are selected. Then for both MarrowTM-10x and MarrowTM-ss2, the genes are scaled to unit variance and only genes with positive mean are selected. Finally, data selected from MarrowTM-10x and MarrowTM-ss2 are concatenated based on the intersection of their gene names.

The MACAQUE-RETINA dataset, which consists of raw macaque single cell count data, as well as its metadata are downloaded from the Single Cell Portal website [20]. As in [18], we only focus on the 30,302 bipolar cells in the total 165,679 cells from macaque retina. There are different levels of batch effect (sample, region and animal) and we only considered the regional batch effect which includes fovea and periphery of retina. As in TM-MARROW, for both the fovea and periphery part of the raw scRNA-seq data, the genes are scaled to unit variance and only genes with positive mean are selected. Then data selected from the fovea and periphery part are concatenated based on the intersection of their gene names.

## Acknowledgements

This research was undertaken using the LIEF HPC-GPGPU Facility hosted at the University of Melbourne. This Facility was established with the assistance of LIEF Grant LE170100200.

## Abbreviations

scRNA-seq: single-cell RNA sequence
scVI: single-cell Variational Inference
MINE: Mutual Information Neural Estimator
MMD: Maximum Mean-Embedding Discrepancy
Pareto MTL: Pareto Multi-Task Learning
ELBO: Evidence Lower Bound
ASW: Averaged Silhouette Width
ARI: Adjusted Rand Index
NMI: Normalized Mutual Information
UCA: Unsupervised Clustering Accuracy
NN: Nearest Neighbor
BE: Batch Entropy

## Availability of data and materials

The datasets analysed during the current study are publicly available, large, and obtained as described in the Single cell RNA-seq datasets section. All the developed code is available in the repository: https://github.com/suswei/single-cell-rna-seq.

## Ethics approval and consent to participate

No ethics approval and consent were required for the study.

## Consent for publication

Not applicable

## Authors’ contributions

Susan Wei led the conceptual design of the study. Hui Li implemented the methodology and performed all experiments. Susan Wei and Hui Li performed the majority of the writing. Heejung Shim and Davis J. McCarthy contributed to supervision of the study. All authors discussed the results and commented on the manuscript. All authors read and approved the final manuscript.

## Additional file 1

Additional file 1 includes Tables S1-S2 and Figures S1-S5.

## Footnotes

1 We represent batch ***s***_*i*_ using dummy encoding, e.g., for B batches, ***s**_i_* ∈ {0,1}^*B*^

2 The estimation of 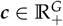 is described in further detail in [6].

3 We use a preliminary version of scVI (version 0.3.0 committed on Mar 6, 2019 on Github), where only ***x*** is the input for encoder.

4 KL divergence (D) between any two probabilities *P* and *Q* are: 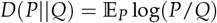.

