## Additional File 1 for "Trade-off between conservation of biological variation and batch effect removal in deep generative modeling for single-cell transcriptomics"

***Table S1.*** **Compare Pareto MTL with MINE and Pareto MTL with MMD.**
The Pareto front–negative ARI/UCA versus negative BE from Pareto MTL with MINE and Pareto MTL with MMD are compared. Better Pareto front has higher percentage/hypervolume/NDC. The values are mean ± standard deviation over 10 Monte Carlos of training-testing split of dataset. Bold illustrates higher mean in comparison.

| Dataset | Percentage | | | | Hypervolume | | | | NDC | | | |
|---|---|---|---|---|---|---|---|---|---|---|---|---|
| | Train | | Test | | Train | | Test | | Train | | Test | |
| | MINE | MMD | MINE | MMD | MINE | MMD | MINE | MMD | MINE | MMD | MINE | MMD |
| **-ARI versus -BE** | | | | | | | | | | | | |
| TM-MARROW | **0.44** ± 0.10 | 0.41 ± 0.13 | **0.49** ± 0.10 | 0.39 ± 0.09 | **0.22** ± 0.01 | 0.13 ± 0.01 | **0.28** ± 0.01 | 0.24 ± 0.01 | **4.8** ± 1.14 | 4.1 ± 1.37 | **5.2** ± 1.03 | 4.4 ± 0.84 |
| MACAQUE-RETINA | **0.37** ± 0.15 | 0.24 ± 0.11 | **0.47** ± 0.19 | 0.34 ± 0.08 | **0.17** ± 0.02 | 0.07 ± 0.01 | **0.16** ± 0.02 | 0.09 ± 0.02 | **3.4** ± 1.26 | 2.6 ± 0.84 | **4.7** ± 1.70 | 3.5 ± 0.85 |
| **-UCA versus -BE** | | | | | | | | | | | | |
| TM-MARROW | **0.47** ± 0.10 | 0.42 ± 0.10 | **0.46** ± 0.10 | 0.40 ± 0.12 | **0.16** ± 0.01 | 0.09 ± 0.01 | **0.22** ± 0.01 | 0.18 ± 0.01 | **5.4** ± 1.26 | 4.4 ± 1.07 | 4.6 ± 0.97 | **4.7** ± 1.25 |
| MACAQUE-RETINA | **0.32** ± 0.16 | 0.27 ± 0.10 | **0.41** ± 0.14 | 0.34 ± 0.12 | **0.13** ± 0.02 | 0.06 ± 0.01 | **0.13** ± 0.02 | 0.07 ± 0.02 | 2.7 ± 1.42 | 2.7 ± 0.67 | **3.4** ± 0.97 | 3.3 ± 1.06 |

***Fig S1.*** **Pareto front in $(\overline{U}_n, \overline{V}_n)$ space estimated via Pareto MTL with MINE.**
We show all 12 Pareto candidates (A) and Pareto front (B) for the generative loss $\overline{U}_n$ versus batch effect measure $\overline{V}_n$ on MACAQUE-RETINA dataset.

**A** **B**

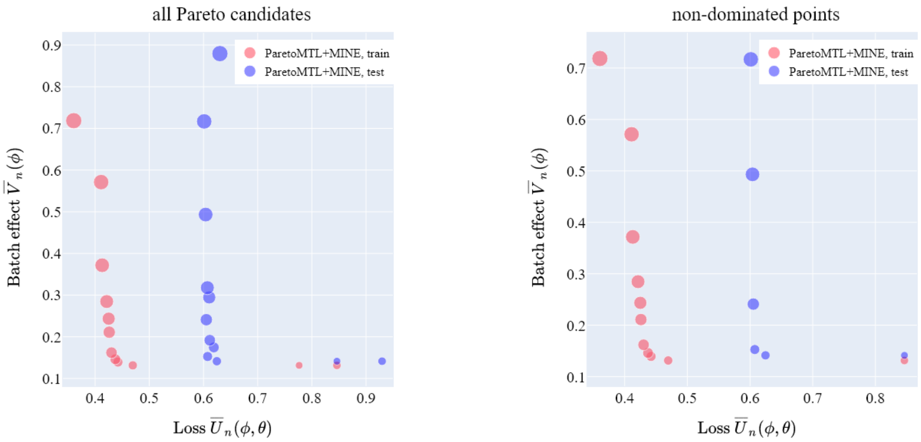

*Fig S2.* **Pareto MTL vs scalarization.**

We show all 12 Pareto candidates (A/C) and Pareto front (B/D) in the $(\overline{U}_n, \overline{V}_n)$ space on MACAQUE-RETINA dataset by Pareto MTL and scalarization when MINE/MMD is used to measure the batch effect. Note that for pareto MTL with MMD and scalarization with MMD, only minimizing $V_n$ does not converge; thus the smaller extreme point does not appear.

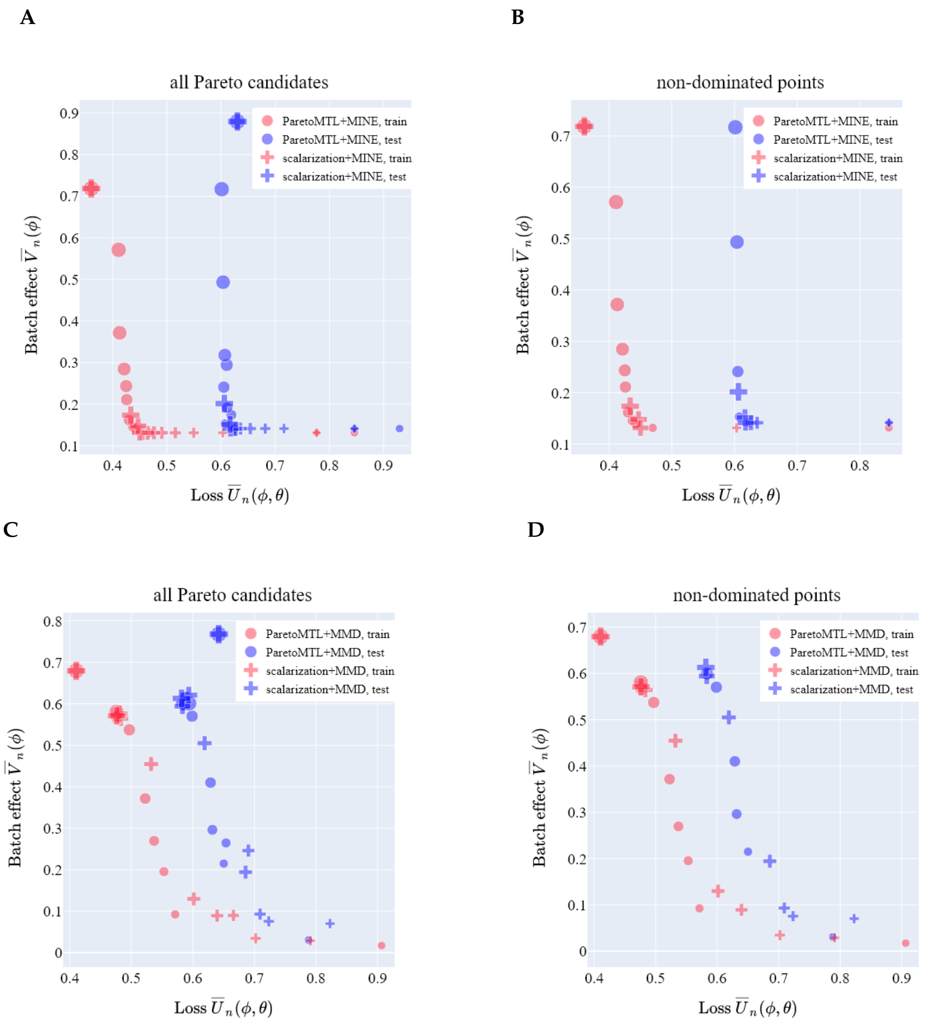


*Fig S3.* **Schemes for hypervolume and NDC.**

Hypervolume is the area of the union of all rectangles determined by the reference point and the estimated non-dominated points (delimited by the red lines in A). For NDC (B), given a set of non-dominated points, suppose $p_a$ and $p_b$ determine the range between maximum and minimum values for the two objectives $L_1$ and $L_2$, and the two ranges can be divided exactly by a pre-specified number $\mu$. The NDC for each square with width $\mu$ is either 0 (no non-dominated points) or 1 (at least one non-dominated points). Sum NDC for all grids to get the NDC of the Pareto front.

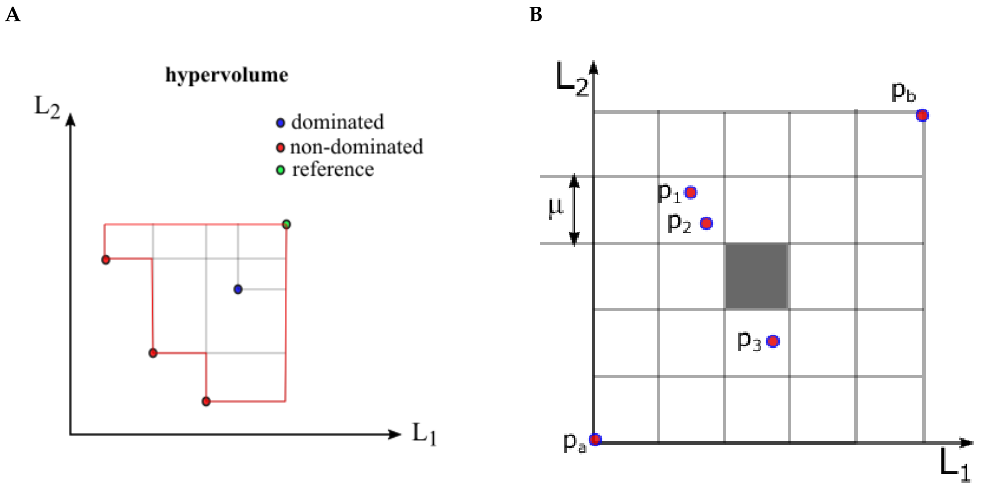


*Fig S4.* **Simulation study 1.**
In order to evaluate MI estimators, we design a simple simulation study where the true MI can be easily calculated as a reference. Suppose $X$ is a categorical variable with 10 categories, and the weight for each category is $p(X = i), i = 0, ..., 9$ s.t. $p(X = i) \geq 0$ and $\sum_{i=0}^{9} p(X = i) = 1$. Suppose $Y$ is a continuous variable, and $p(Y|X = i) \sim \mathrm{N}(\mu_i \mathbf{1}_2, \sigma_i^2 \boldsymbol{I}_2)$, where $i = 0, ..., 9$, $\mathbf{1}_2$ is a two-dimensional vector with 1 as its elements, and $\boldsymbol{I}_2$ is a 2-by-2 identity matrix. By varying $p(X = i)$, $\mu_i$ and $\sigma_i$, we design 7 cases. For each case, the true MI and the MINE estimator are calculated. Besides, the empirical (i.e. sample average of the log ratio of the joint probability and the product of marginal probabilities) and NN (i.e. Nearest Neighbor) estimators of the true MI are also provided. We evaluate the empirical and NN estimators since they can be used as reference when true MI is intractable with known distributions (use empirical estimator as reference) or unknown distributions (use NN estimator as reference). The true MI for the first case is 0. Overall, MINE, empirical and NN estimators approximate the true MI well. However, the caveat is that the empirical and NN estimators are not suitable for deep learning training.

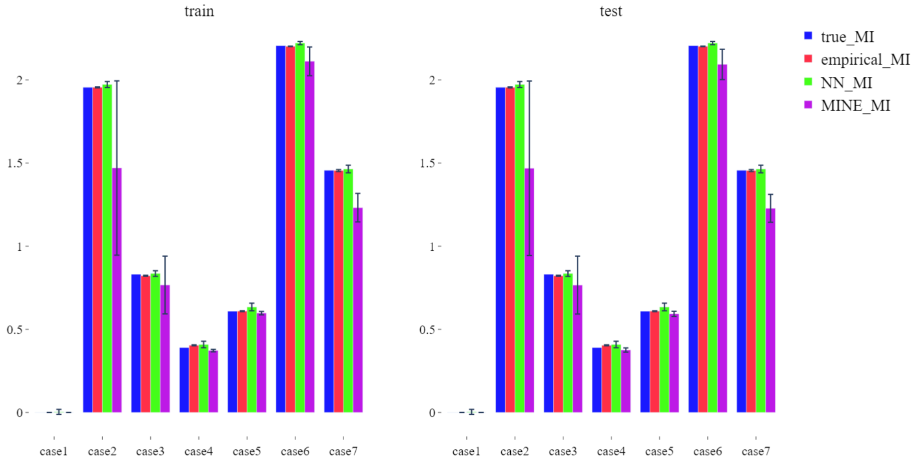

*Fig S5.* **Simulation study 2.**
In order to evaluate MI estimators in complex situations, we design a simulation study where the true MI is hard to calculate and the empirical estimator is used as reference since the probabilities are known. Suppose $X$ is a categorical variable with 2 categories, and the weight $p(X=i)=0.5, \forall i=0,1$. Suppose $Y$ is a continuous variable with dimension $d$, and $P(Y|X=i), \forall i=0,1$ is a Gaussian mixture with $m$ components. Specifically, $p(Y|X=0)=\sum_{j=1}^{m}(1/m)N(\mu_j,\sigma_j^2\boldsymbol{I}_d)$, where $\mu_j$ is randomly sampled from standard Gaussian with dimension $d$, and $\sigma_j=1$. In contrast, $p(Y|X=1)=\sum_{j=1}^{m}(1/m)N(\mu_j,\sigma_j^2\boldsymbol{I}_d)$, where $\mu_j$ is randomly sampled from $N(\mu\mathbf{1}_d,\boldsymbol{I}_d)$, $\mu$ can be any real number, $\mathbf{1}_d$ is a $d$-dimensional vector with 1 as its elements, $\boldsymbol{I}_d$ is a $d$-by-$d$ identity matrix, and $\sigma_j=1$ or $\sigma_j=0.9\mu$. By varying $d$, $m$, $\mu$, and by choosing from $\sigma_j=1$ or $\sigma_j=0.9\mu$ for $p(Y|X=1)$, 16 cases are designed. With the empirical estimate for the true MI as reference, we conclude that MINE and NN are good MI estimators between a continuous and discrete random variable in complex situations.

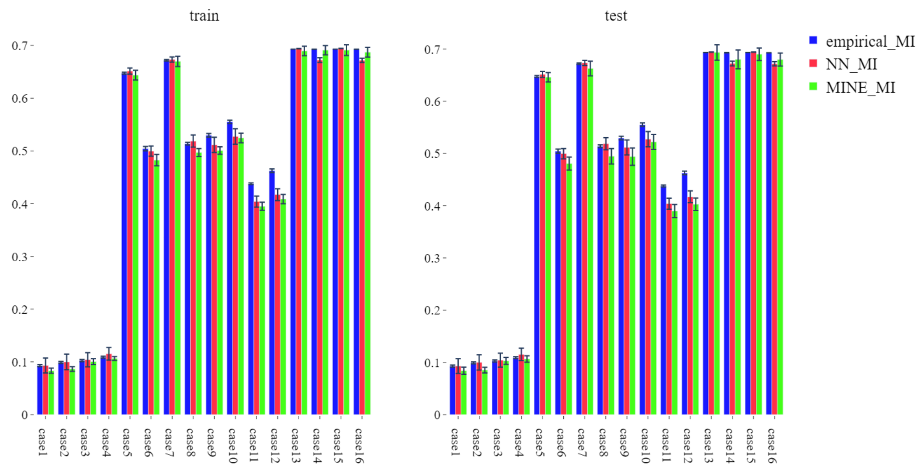

***Fig S6.*** **The trade-off curves of surrogate metrics.**
Pareto MTL with MINE and Pareto MTL with MMD are compared for the Pareto candidates and Pareto front—negative ARI/UCA versus negative BE on the TM-MARROW dataset.

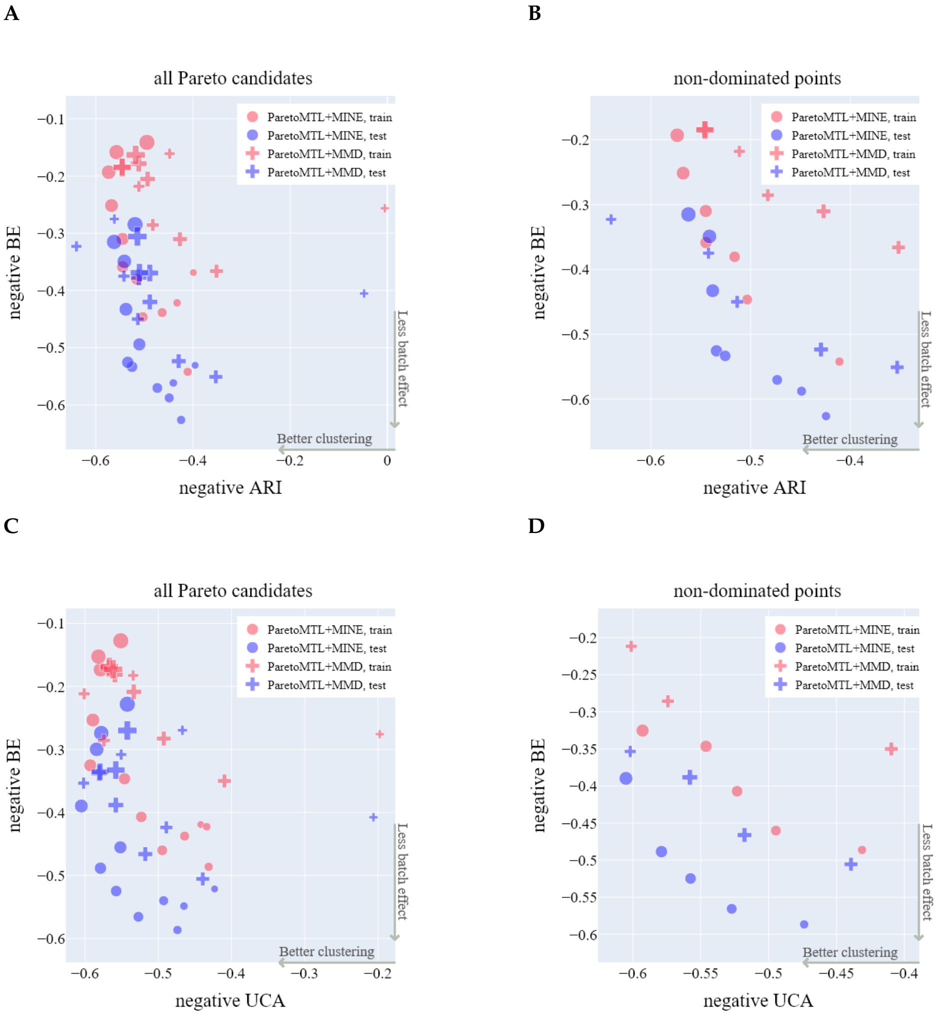

***Fig S7.*** **The trade-off curves of surrogate metrics between scalarization with MINE and scalarization with MMD.**

We show the trade-off curves (both all Pareto candidates and Pareto front) for $U_n$ versus $NN_n$ and negatve ASW/NMI versus negative BE between scalarization with MINE and scalarization with MMD on TM-MARROW dataset.

**A**

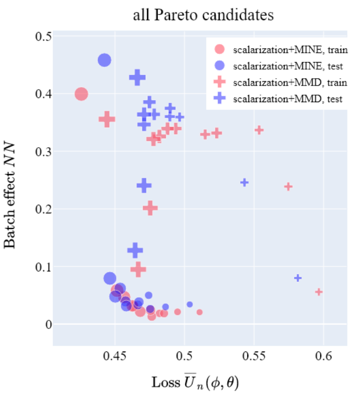


**B**

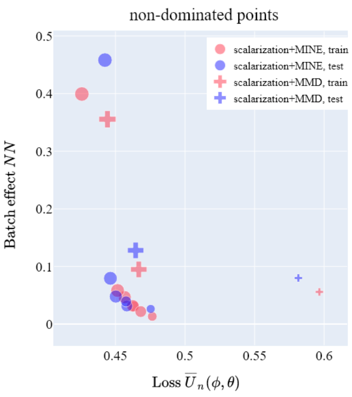


**C**

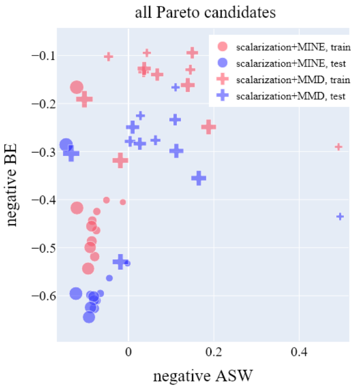


**D**

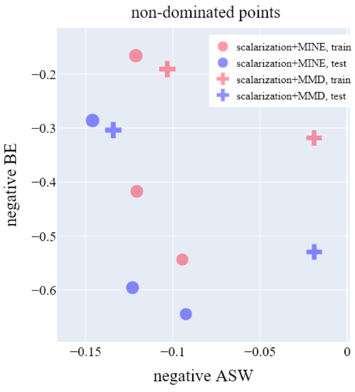


**E**

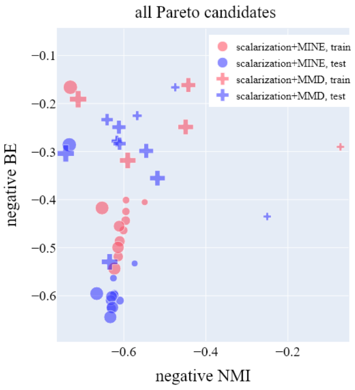


**F**

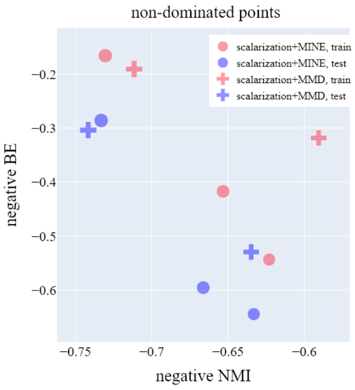
